# Insights from the genomes of four diploid *Camelina* spp.

**DOI:** 10.1101/2021.08.23.455123

**Authors:** Sara L. Martin, Beatriz Lujan Toro, Tracey James, Connie A. Sauder, Martin Laforest

## Abstract

Plant evolution has been a complex process involving hybridization and polyploidization making understanding the origin and evolution of a plant’s genome challenging even once a published genome is available. The oilseed crop, *Camelina sativa* (*Brassicaceae)*, has a fully sequenced allohexaploid genome with three unknown ancestors. To better understand which extant species best represent the ancestral genomes that contributed to *C. sativa*’s formation, we sequenced and assembled chromosome level draft genomes for four diploid members of *Camelina*: *C. neglecta C. hispida* var. *hispida*, *C. hispida* var. *grandiflora* and C. *laxa* using long and short read data scaffolded with proximity data. We then conducted phylogenetic analyses on regions of synteny and on genes described for *Arabidopsis thaliana,* from across each nuclear genome and the chloroplasts to examine evolutionary relationships within *Camelina* and *Camelineae*. We conclude that *C. neglecta* is closely related to *C. sativa*’s sub-genome 1 and that *C. hispida* var. *hispida* and *C. hispida* var. *grandiflora* are most closely related to *C. sativa*’s sub-genome 3. Further, the abundance and density of transposable elements, specifically *Helitrons*, suggest that the progenitor genome that contributed *C. sativa*’s sub-genome 3 maybe more similar to the genome of *C. hispida* var. *hispida* than that of *C. hispida* var. *grandiflora.* These diploid genomes show few structural differences when compared to *C. sativa*’s genome indicating little change to chromosome structure following allopolyploidization. This work also indicates that *C. neglecta* and *C. hispida* are important resources for understanding the genetics of *C. sativa* and potential resources for crop improvement.

## Introduction

A key goal in evolutionary biology is understanding the evolution of genomes - how changes in structure and content affects the rate of evolution (Otto and Whitton 2002). Three linked processes shape genome structure: hybridization (Stebbins 1968; Rieseberg 1997; Soltis and Soltis 2009; Abbott et al. 2013), polyploidization (de Wet 1971; Levin 1983; Soltis and Soltis 1999; Husband et al. 2013) and chromosomal rearrangements (Rieseberg 2001; Rieseberg and Willis 2007). Crop genomes such as maize, canola, soybean, sugarcane and wheat (Schmutz et al. 2010; Schnable et al. 2011; Chalhoub et al. 2014; The International Wheat Genome Sequencing Consortium 2014) have underscored the frequency of these processes. While less effort has focused on diploid wild crop relatives (Michael and VanBuren 2015), sequencing the extant representatives of crop progenitors can provide insight into the crop’s origins and genomic evolution (Marcussen et al. 2014; Latta et al. 2019).

The oil seed crop and allohexaploid *Camelina sativa* (L.) Crantz (2n = 40) has been sequenced and well described (Kagale et al. 2014). It’s three parental genomes are thought to have diverged from each other recently between 2.5 Mya (Žerdoner Čalasan et al. 2019) and 5.4 Mya (Kagale et al. 2014) with the dates of the hybridizations contributing to *Camelina sativa*’s formation estimated as 5-10,000 years ago (Kagale et al. 2014). The *Camelina* genus includes four taxa with known diploid chromosome counts: *Camelina neglecta* (2n = 12, 1C = 265 Mb) J. Brock, Mandáková, Lysak & Al-Shehbaz, *Camelina laxa* C. A. Mey. (2n = 12, 1C = 275 Mb), *Camelina hispida* Boiss. var. *hispida* (2n = 14, 1C = 355 Mb) and *Camelina hispida* var*. grandiflora* (Boiss.) Hedge (2n = 14, 1C = 315 Mb) (Al-Shehbaz 2012; Martin et al. 2017; Brock et al. 2019). These species could be modern representatives of genomes involved in *C. sativa*’s formation.

While determining the origin of a polyploid lineage is generally difficult (Kyriakidou et al. 2018; Rothfels 2021), our knowledge of the ancestral chromosome structure of the Camelineae provides an opportunity to investigate related diploid genomes. In the 2000’s, researchers defined 24 large conserved collinear regions or blocks (Ancestral Crucifer Karyotype or ACK) among crucifer genomes (Schranz et al. 2006; Murat et al. 2015; Lysak et al. 2016) and reconstructed an ancestral karyotype with 8 chromosomes similar to *Arabidopsis lyrata* (L.) O’Kane & Al-Shehbaz (Koch and Kiefer 2005). These blocks were later grouped into 16 ancestral karyotype regions (ABK) corresponding to ancestral chromosome arms (Murat et al. 2015). *Camelina sativa*’s sub-genomes each show a conserved ACK structure, with 21 in-block breaks that, most likely, occurred in its progenitors (Lysak et al. 2016). Here we extend our understanding of the evolution of *C. sativa*by assembling draft genomes for *C. neglecta*, *C. laxa, C. hispida* var. *hispida* and *C. hispida* var. *grandiflora.* Using this data and the ACK structure we phase *C. sativa*’s sub-genomes and examine relationships within the Camelineae using a phylogenetic analysis before delving further into *C. sativa’s* potential paternal lineage using transposable element abundance.

## Materials and Methods

### Plant Material and Nucleic Acid Isolation

*Camelina neglecta, C. hispida var. hispida, C. hispida var. grandiflora,* and *C. laxa* seed were obtained from the North Central Regional Plant Introduction Station (NCRPIS) (PI650135, PI650139, PI650133 and PI633185, respectively). Seeds obtained from NCRPIS were stratified in petri dishes using filter paper that was moistened with 0.2% KNO_3_, sealed with Parafilm (Pechiney Plastic Packaging Company, Illinois, USA), and placed at 4°C in the dark for 2 weeks. For seed germination, the plates were then placed at room temperature under growth lights with a 16 h/8 h day/night light cycle. Seedlings were then sown on soil (soil, peat, and sand; 1:2:1; Promix, Rivière-du-Loup, Québec, Canada) with a photoperiod of 16 h 20°C days/8 h 18°C nights. For the largely self-incompatible species, *C. hispida* and *C. laxa,* rosette leaves were sampled using dry ice for DNA extractions and stored at -80. For the self-compatible, *C. neglecta*, after 6 weeks, the plants were placed for another 6 weeks at 4°C with an 8 h/16 h day/night photoperiod for vernalization. Plants were then transplanted to 5 in pots with the same soil and allowed to self-pollinate and set seed in a 16 h/8 h day/night photoperiod with 20°C days and 18°C nights. Following approximately 3 months, the mature seeds were collected and this process was repeated to obtain a fifth-generation inbred line before sampling and storage. Vouchers for each of the accessions used have been deposited in the DAO (Department of Agriculture Ottawa) herbarium (*C. neglecta* DAO 902176; *C. laxa*: DAO 902754; *C. hispida* var. *hispida*: DAO 902780; and *C. hispida* var. *grandiflora:* DAO 902768).

For Pacific Biosciences long read (PacBio; Pacific Biosciences, Menlo Park, CA, USA) and Illumina short reads (Illumina Inc., San Diego, California, U.S) sequencing, total DNA was extracted using a FastDNA Spin Kit (MP Biomedicals, Solon, OH), grinding was done in the FastPrep (MP Biomedicals) at 4.0 for 20 s, with the addition of one ceramic bead. Two DNA extractions were pooled for a total volume of 200 μl and precipitated with the addition of 20 μl 3 M NaOAc and 200 μl 100% ethanol. Following overnight incubation at -20°C, the DNA was centrifuged at 13,000 rpm at 4°C for 30 min. The ethanol was decanted and the DNA pellet was washed with cold 70% ethanol, dried at 37°C for approximately 20 min and re-suspended in 100 μl 5 mM Tris-HCl (pH 8.5). The DNA concentration was determined by Qubit was 110 ng/μl. DNA quality was determined by running 1.0 μl on a 0.8% E-gel (Invitrogen by ThermoFisher) beside a 0.2 μg of 20kb ladder (GeneRuler 1kbPlus, ThermoFisher).

DNA was also extracted to generate sequence data using Oxford Nanopore Technologies (ONT; Oxford Nanopore Technologies, Oxford Science Park, UK). High molecular weight DNA extraction procedures were carried out as described in Workman et al. (Workman et al. 2018). The Short Read Eliminator kit (Circulomics Inc) was used as per the directions.

### Sequencing

Sequencing was conducted in five different locations. McGill University’s Genome Quebec Innovation Centre completed PacBio sequencing using P6-C4 chemistry for *C. laxa, C. hispida* var. *hispida* and *C. neglecta*. A total of 7 Single Molecule Real-Time (SMRT) cells were used for sequencing *C. neglecta* and 8 cells each were used for *C. laxa* and *C. hispida*. They also generated paired end Illumina data for *C. laxa*, *C. hispida* var. *hispida* and *C. hispida* var. *grandiflora* with runs of 2 x 150 bases. The sequencing facility at the Microbial Molecular Technologies Laboratory (MMTL) in Ottawa (Ottawa Research and Development Centre, Agriculture and Agri-Food Canada) was used for paired-end sequencing using Illumina MiSeq v3 chemistry with runs of 2 x 300 bases with 500bp inserts. The Centre for Applied Genomics, The Hospital for Sick Children, Toronto, Canada prepared and sequenced four additional libraries: one paired-end (PE) library with 550 bp inserts using a Nano kit (Illumina, San Diego, CA, USA) and three mate-pair (MP) libraries: 3kb, 5kb and 10kb inserts using the Nextera mate-pair kit (Illumina). All four libraries were sequenced on an Illumina HiSeq-2000 using v4 chemistry and flow cells with runs of 2 × 126 bases. ONT sequencing data was obtained using a MinION with one FLO-MIN106 flow cell run for 48h for each taxon in the Martin Laboratory. Base pair calling was completed with Guppy v. 3.2.2+9fe0a78 (Wick et al. 2019). Finally, library preparation for chromosome conformation capture (Hi-C) analyses used the Proximo Hi-C 2.0 Kit from Phase Genomics Inc. (Seattle, WA, USA) and sequenced using Ilumina HiSeq 4000.

### Quality Control and *de novo* Genome Assembly

Raw PacBio and ONT reads were self-corrected and assembled using Canu v1.8 (Koren et al. 2016). The corrected error rate was set to 14.4% for both *C. hispida* var. *grandiflora*, where we only had ONT data, and for *C. hispida* var. *hispida*. Draft assemblies were polished using Illumina data filtered and trimmed with trimmomatic v 0.33 (Bolger et al. 2014) and up to three iterations of Pilon v 1.23 (Walker et al. 2014) using bowtie2 v 2.3.4.3 (Langmead and Salzberg 2012) and Burrows-Wheeler Aligner (bwa) (Li 2013). For C. *hispida* and *C. laxa*, the program Purge Haplotigs (Roach et al. 2018) was used to reduce redundancy resulting from the heterozygosity.

The quality of the polished assemblies was evaluated with QUAST 5.0.2 (Gurevich et al., 2013). The completion of the assembly’s gene space was evaluated using BUSCO v3.0.2 (Simão et al. 2015) using the embryophyta_odb10 database and the degree to which the assemblies had been successfully collapsed was further assessed with HapPy (Guiglielmoni et al. 2021). Final heterozygosity and error rates were calculated with samtools v 1.9 (Li et al. 2009) and SNP detection and phasing was completed with Princess v0.01 (Mahmoud et al. 2021).

The chloroplast genome for each species was extracted from the draft assemblies by aligning contigs with *C. sativa*’s chloroplast genome using nucmer v 4.0 from MUMmer v 4.0. Chloroplast contigs were ordered and oriented based on *C. sativa*’s chloroplast and merged into a consensus using a custom script written in R.

### Genome Scaffolding

Phase genomics used their proprietary software, Proximo, to produce chromosome level scaffolds (Oddes et al. 2018) for *C. neglecta*, *C. hispida* var. *hispida* and *C. laxa*. The scaffolding tool, ntJoin 1.0.3-0 (Coombe et al. 2020), was used to scaffold the genome of *C. hispida* var. *grandiflora*using the Hi-C scaffolded assembly of *C. hispida* var. *hispida*. A final round of polishing by Pilon was completed following scaffolding.

All genomes are available from as part of Bioproject PRJNA750147.

### Phylogenetic analysis of *C. sativa s*ub-genomes and Camelineae diploids

We determined the phylogenetic relationships among genomes for 1) diploid members of the Camelineae: *Arabidopsis lyrata* (Ensembl Genomes version 1.0), *Capsella rubella* (NCBI v. ANNY00000000.1), *Neslia paniculata* (L.) Desvaux (S. Wright personal communication) 2) the three sub-genomes of *C. sativa* (NCBI version JFZQ0000000.1) and 3) the four diploid *Camelina* species sequenced here. We used three methods to extract regions of the genomes for analysis. First we identified random fragments within homologous regions of the genomes, second we used a reciprocal best hit method to identify shared genes from *Arabidopsis thaliana* (TAIR version 10), and third we used Orthofinder 2.3.11 (Emms and Kelly 2017; Emms and Kelly 2019).

For the first approach shared homologous regions within each ACK block as described in the *A. lyrata* genome (Schranz et al. 2006) were isolated (Table S1). Each ACK region was cut into 1000 bp fragments, aligned to *A. lyrata*’s genome using bowtie2 (Langmead and Salzberg 2012) and filtered. Pairwise collinearity between this set of filtered fragments and each taxon was determined using nucmer from MUMmer (v4.0) then a custom R script determined which fragments that overlapped between all genomes and were found in three copies within a consistent set of *C. sativa*’s chromsomes (Table S2). Each set was aligned using msa 1.18.0 (Bodenhofer et al. 2015) and run in MrBayes v3.2.1 using a stepping stone model run for 6,000,000 generations to determine the appropriate model (Ronquist et al. 2012). We then ran the preferred model for 5,000,000 generations checking convergence of each run with the r package rwty (Warren et al. 2017). We then calculated seven metrics to exclude biased sequences or sources of misleading phylogenetic signal using TreeCmp 2.0 (Bogdanowicz et al. 2012) and TreSpEx 1.1 (Struck 2014). Specifically, following Nikolov et al. (2019) we calculated the number of matching splits and the Robinson-Foulds tree distances using TreeCmp; the upper quartile and standard deviation of the long-branch scores, average patristic differences, and R^2^ of the saturation score and slope with TreSpEx. Fragments were excluded from further consideration if they failed the convergence checks or were outliers for one or more of the seven phylogenetic metrics at the 99^th^ percentile. For each set of trees belonging to ABKs located on the same chromosome or chromosome arm, we estimated species trees with ASTRAL-III (5.1.1) (Zhang et al. 2018).

Our second method used the information phasing the sub-genome structure of *C. sativa* as determined by the fragments above, to identify reciprocal best hits (RBH) (Chen et al. 2017) for *A. thaliana* genes in each genome and sub-genome. Specifically, sequences for genes identified for *A. thaliana* in the TAIR 10 assembly (available from www.arabidopsis.org) were used to extract similar sequences for each genome. Following recommendations by Chen et al. (2017) BLAST 2.2.31 (Altschul et al. 1990) hits for *A. thaliana* genes were screened (e-value of less or equal to 0.0001, 70% or more of the query length, 70% or greater identity with query, a bit score ratio between the first and second BLAST hit of 1.2 or greater) extracted from each target genome and BLASTed back to *A. thaliana*’s genome and filtered again with the additional criteria of aligning to the gene’s original location.

The phylogenetic analysis of these best hits was then completed as above using MrBayes and ASTRAL-III by ABK. In addition, we randomly selected one sequence from each ACK 25 times to be a set of unlinked data and analyzed these using StarBEAST2 2.6.3 (Ogilvie et al. 2017; Suchard et al. 2018; Bouckaert et al. 2019), PhyloNet 3.8.2 (Than et al. 2008; Wen et al. 2018; Cao et al. 2019) and to generate a consensus tree with ASTRAL-III. StarBeast2 was run for between 20 and 100 million generations as required to produce estimated sample sizes (ESS) values above 200 with the GTR site model using a configuration file created by BEAUti 2.6.3 (Bouckaert et al. 2019), convergence was examined with Tracer (1.7.1) (Rambaut et al. 2018), summarized with TreeAnnotator 2.6.3 (Bouckaert et al. 2019) and plotted with the densiTree function in the R package phangorn 2.5.5 (Schliep 2011). PhyloNet’s MCMC_Seq command was run with chain lengths of between 20 and 80 million as required to produce ESS values above 200 and the maximum number of reticulations set to 4. Networks and trees produced by PhyloNet were visualized using plotTree function from the package phytools 0.7.20 (Revell 2012).

Finally, AUGUSTUS 3.3.2 (Stanke et al. 2006) was run on the final version of the assembled genomes, reference genomes, and *Camelina sativa*’s phased sub-genomes to predict genes *in silico*using *Arabidopsis*. Amino acid sequences were then provided to OrthoFinder (Emms and Kelly 2019), which estimated the species tree.

Entire chloroplast genomes for *Arabis alpina*, *Arabidopsis thaliana*, *A. lyrata*, *Capsella bursa-pastoris*, *C. grandiflora*, *C. rubella* and *Camelina sativa* were downloaded from the Chloroplast Genome Database (https://rocaplab.ocean.washington.edu/old_website/tools/cpbase) and aligned with the four *Camelina* chloroplasts using the msa function in R (see above). A consensus tree was then estimated with MrBayes as above. Annotation of the chloroplast genomes was completed with GeSeq 1.77 (Tillich et al. 2017) through CHLOROBOX website (https://chlorobox.mpimp-golm.mpg.de/index.html), using *Camelina sativa*, *A. lyrata,* and *Capsella rubella*’s chloroplasts as reference sequences. The chloroplast genomes with their annotations were then visualized with the website’s OrganellarGenomeDRAW (OGDRAW) 1.3.1 (Greiner et al. 2019).

### Synteny between Camelina diploids and C. sativa and A. lyrata

To evaluate collinearity between the chromosome level draft genomes we used nucmer with both *C. sativa* and *A. lyrata* as references for comparison and scripts written in R employing the package circlize 0.4.11 (Gu et al. 2014) to order and visualize the alignments.

### Transposable Element Annotation

Transposable elements (TEs) were located using the Extensive *de novo* TE Annotator 1.8.3 (EDTA; Ou et al., 2010), EAHelitron 1.5.1 (Hu et al. 2019), LRT_FINDER/LTR_FINDER_parallel (Xu and Wang 2007; Ou and Jiang 2019) and LTRharvest (Ellinghaus et al. 2008). Data from these last two programs was processed with LRT_retreiver (Ou and Jiang 2018), which was used to calculate the LTR Assembly Index, (LAI) (Ou et al. 2018). We determined whether *Helitrons* detected in *C. sativa*’s sub-genome 3 were also detected in syntenic regions of *C. hispida var. hispida* and *C. hispida* var. *grandiflora* using a script written in R and output from EAHelitron and nucmer. Specifically, we examined all regions of synteny (determined by nucmer), determined if a *Helitron* was detected in the region (by EAHelitron using the presence of sequence CTAG and a GC rich hairpin), and examined sequence similarity for 1000 bp upstream of the 3’ end. We considered any pairs with 80% identity to represents the same *Helitron*insertion event.

### Additional software used

The version of R used was 3.6.3 (2020-02-29) -- “Holding the Windsock.” Sequence handling, tree plotting and graphical display were facilitated by numerous R packages in addition to those mentioned above including: ape 5.0 (Paradis and Schliep 2019), apex 1.0.4 (Schliep et al. 2020), Biostrings 2.56.0 (Pagès et al. 2020), pals 1.7 (Wright 2021), pBrackets 1.0.1 (Schulz 2021), plotrix 3.7.8 (Lemon 2006), plyr 1.8.6 (Wickham 2011), ips 0.0.11 (Heibl 2008), IRanges 2.22.2 (Lawrence et al. 2013), outliers 0.14 (Komsta 2011), RIdeogram 0.2.2 (Hao et al. 2020), Rsamtools 2.4.0 (Morgan et al. 2020), seqinr 3.6.1 (Bastolla et al. 2007), stringr 1.4.0 (Wickham 2019), treeio 1.10.0 (Wang et al. 2020), and VennDiagram.1.7.1 (Chen 2021).

The program FigTree 1.4.4 (Rambaut 2018) was used to convert trees including their support values to a format easily readable in R.

Unless otherwise specified all tools were run using default settings.

## Results

### Genome Assemblies

Sequencing coverage of the genome differed for each species with *C. neglecta* receiving the majority of our sequencing efforts (286x raw coverage) (Table 1). Following assembly with Canu and polishing with Pilon, *C. neglecta* had the most contiguous assembly with 204 contigs and an NG50 of 11,493,634 (Table 2). Scaffolding using Hi-C data and Phase Genomics’ Proximo resulted in chromosome level assemblies for *C. neglecta* (n = 6), *C. hispida* var. *hispida* (n = 7) and *C. laxa* (n =6) with approximately 70% or more of their expected lengths and NG50s of 29,279,412. 39,460,631, and 31,147,072 respectively. Following scaffolding by ntJoin, 70% of the expected genome length for *C. hispida* var. *grandiflora* was incorporated into a chromosome level assembly (n = 7). The completeness of gene space, as evaluated by estimating the proportion of core conserved eukaryotic genes recovered by the Benchmarking Universal Single Copy Orthologs (BUSCO) score, indicated that all assemblies had at least 90% of the expected genes (Table 2). The small percentage of duplicated BUSCO genes indicated that the assemblies have largely been collapsed. This was confirmed by haploidy scores from HapPy of over 0.9. The genomes all rank as gold quality based the high level of intact LTR elements detected in the genome (LIA > 20) (Table 2).

**Table 1.** Genome sizes (1C) as estimated from flow cytometry and the fold coverage (x) for the four genomes sequenced here provided by long read data (PacBio and ONT) and short read data including Illumina paired-end reads (PE) and Illumina mate pairs (MP) with upper limits of estimates of heterozygosity (H) and error rates (ER) calculated by GenomeScope from the Illumina data.

| SPECIES | GENOME<br>SIZE (MB) | PACBIO | ONT | ILLUMINA |  | RAW<br>COVERAGE | GENOMESCOPE |  |
| --- | --- | --- | --- | --- | --- | --- | --- | --- |
|  |  |  |  | PE | MP |  | H | ER |
| *Camelina neglecta* | 265 | 38 | 60 | 92 | 96 | 286 | 0.17% | 0.08% |
| *Camelina hispida* var. *hispida* | 355 | 38 | 53 | 52 | - | 143 | 1.09% | 0.15% |
| *Camelina hispida* var. *grandiflora* | 315 | - | 32 | 64 | - | 96 | 2.14% | 0.15% |
| *Camelina laxa* | 275 | 57 | 74 | 47 | - | 177 | 1.36% | 0.24% |

**Table 2.** Statistics on genome assemblies for *C. neglecta*, *C. hispida* var. *hispida*, *C. hispida* var. *grandiflora*, and *C. laxa* generated by QUAST and BUSCO based on the embryophyta_odb10 database before and after scaffolding. The initial assembly values for *C. hispida* and *C. laxa* are for assemblies following assembly by Canu and reduction in the number of alternative contigs resulting from the heterozygosity of these genomes by Purge Haplotigs. All assemblies were polished with Pilon. Addition metrics are 1) the haploidy score (HapPy) with 1 indicating perfect collapse, 2) the LTR Assembly Index (LAI) (LTR_retriever) with scores > 20 indicating gold quality, 3) final error rate (samtools), 4) number of SNPs (PRINCESS) and 5) final heterozygosity (PRINCESS).

|  | *Camelina neglecta* |  | *Camelina hispida* var. *hispida* |  | *Camelina hispida* var. *grandiflora* |  | *Camelina laxa* |  |
| --- | --- | --- | --- | --- | --- | --- | --- | --- |
| ASSEMBLY STRATEGY | CANU | HI-C | CANU & PURGE HAPLOTIGS | HI-C | CANU & PURGE HAPLOTIGS | NTJOIN | CANU & PURGE HAPLOTIGS | HI-C |
| contigs (>= 0 bp) | 204 | 6 | 747 | 7 | 1,072 | 7 | 900 | 6 |
| contigs (>= 10000 bp) | 157 | 6 | 703 | 7 | 896 | 7 | 785 | 6 |
| contigs (>= 50000 bp) | 139 | 6 | 552 | 7 | 748 | 7 | 530 | 6 |
| Total length (>= 0 bp) | 210,252,877 | 194,776,448 | 305,512,316 | 283,171,403 | 247,512,955 | 247,668,533 | 223,800,267 | 199,991,310 |
| Total length (>= 10000 bp) | 210,014,359 | 194,776,448 | 305,328,622 | 283,171,403 | 247,070,138 | 247,668,533 | 223,546,568 | 199,991,310 |
| Total length (>= 50000 bp) | 209,462,969 | 194,776,448 | 301,174,771 | 283,171,403 | 242,642,354 | 247,668,533 | 217,296,729 | 199,991,310 |
| Largest contig | 19,486,679 | 48,079,882 | 6,147,326 | 45,593,769 | 3,518,166 | 41,100,942 | 4,931,297 | 39,604,730 |
| Total length | 210,252,877 | 194,726,046 | 305512316 | 283,171,403 | 247,512,955 | 247,668,533 | 223,800,267 | 199,991,310 |
| Estimated genome size | 265,000,000 | 265,000,000 | 360,000,000 | 360,000,000 | 320,000,000 | 320,000,000 | 275,000,000 | 275,000,000 |
| Percent of Expected | 79% | 74% | 85% | 79% | 77% | 77% | 81% | 73% |
| N50 | 13,701,387 | 30,499,177 | 1053466 | 41,586,011 | 477,640 | 37,976,760 | 651,797 | 31,824,321 |
| NG50 | 11,493,634 | 29,279,412 | 843316 | 39,460,631 | 346,627 | 30,680,265 | 477,199 | 31,147,072 |
| N's per 100 kbp | - | 1 | - | 16 | - | 32,233 | - | 20 |
| BUSCO | 2,121 | 2,121 | 2,121 | 2,121 | 2,121 | 2,121 | 2,121 | 2,121 |
| Complete (%) | 2,082 (98) | 2,084 (98) | 2055 (97) | 2,025 (96) | 2,063 (97) | 1,901 (90) | 2,079 (98) | 2032 (96) |
| Complete Single Copy (%) | 2,051 (97) | 2,055 (97) | 1938 (91) | 1,971 (93) | 1,863 (88) | 1,853 (88) | 1,939 (91) | 1,978 (93) |
| Complete Duplicated (%) | 31 (1) | 29 (1) | 117 (6) | 54 (3) | 200 (9) | 48 (2) | 140 (7) | 54 (3) |
| Fragmented (%) | 14 (<1) | 14 (<1) | 28 (1) | 43 (2) | 18 (<1) | 65 (3) | 14 (<1) | 18 (<1) |
| Missing (%) | 25 (1) | 23 (1) | 38 (2) | 53 (3) | 40 (2) | 155 (7) | 28 (1) | 71 (3) |
| Haploidy Score |  | 0.99 |  | 0.97 |  | 0.96 |  | 0.94 |
| LAI |  | 21.1 |  | 25.7 |  | 27.6 |  | 22.0 |
| Error Rate |  | 0.004 |  | 0.007 |  | 0.02 |  | 0.007 |
| SNPs |  | 420,396 |  | 1,564,682 |  | 1,813,558 |  | 1,488,895 |
| Heterozygosity |  | 0.002 |  | 0.006 |  | 0.007 |  | 0.008 |

### Phylogenetic relationships among *C. sativa* and Camelineae diploids

In total 1,444 fragments within the ACK blocks described for *A. lyrata* genome were found to be shared among the eight core Camelineae taxa with three copies distributed across *C. sativa*’s genome. Fragments from each ancestral chromosome showed the same pattern of localization in *C. sativa*’s genome in the majority of cases (Table 3). For example ACK blocks A, B, and C from ancestral chromosome 1 all showed the highest number of hits on *C. sativa*’s chromosomes 3, 14 and 17. Where differences occurred, there was consistency corresponding to ancestral chromosome arms and, therefore, ABK group. The pattern of hits across the genomes resulted in eleven groupings, corresponding to either ancestral chromosome arms or entire chromosomes, that were used in further phylogenetic analyses (Table 4).

**Table 3.** Distribution of the top three chromosomes with successful alignments of fragments from each of *Arabidopsis lyrata’s* ACKs across the genome of *Camelina sativa* as determined by bowtie2 ordered by number of hits.

| ACK | ABK | *Camelina sativa* TOP<br>CHROMOSOME | *Camelina sativa* 2<sup>nd</sup><br>CHROMOSOME | *Camelina sativa* 3<sup>rd</sup><br>CHROMOSOME |
| --- | --- | --- | --- | --- |
| A | AK2 | 3 | 14 | 17 |
| B | AK2 | 14 | 3 | 17 |
| C | AK1 | 3 | 14 | 17 |
| D | AK3 | 7 | 16 | 9 |
| E | AK4 | 7 | 16 | 9 |
| F | AK6 | 1 | 15 | 19 |
| G | AK6 | 1 | 15 | 19 |
| H | AK5 | 15 | 1 | 19 |
| I | AK7 | 7 | 16 | 9 |
| J | AK8 | 5 | 6 | 4 |
| K | AK9 | 4 | 6 | 9 |
| L | AK9 | 4 | 6 | 9 |
| M | AK10 | 9 | 6 | 4 |
| N | AK10 | 4 | 6 | 9 |
| O | AK11 | 2 | 13 | 8 |
| P | AK11 | 13 | 2 | 8 |
| Q | AK12 | 8 | 13 | 20 |
| R | AK12 | 8 | 13 | 20 |
| S | AK13 | 10 | 11 | 12 |
| T | AK14 | 10 | 11 | 12 |
| U | AK14 | 11 | 12 | 10 |
| V | AK15 | 11 | 20 | 18 |
| W | AK16 | 18 | 11 | 2 |
| X | AK16 | 11 | 2 | 18 |

**Table 4.**
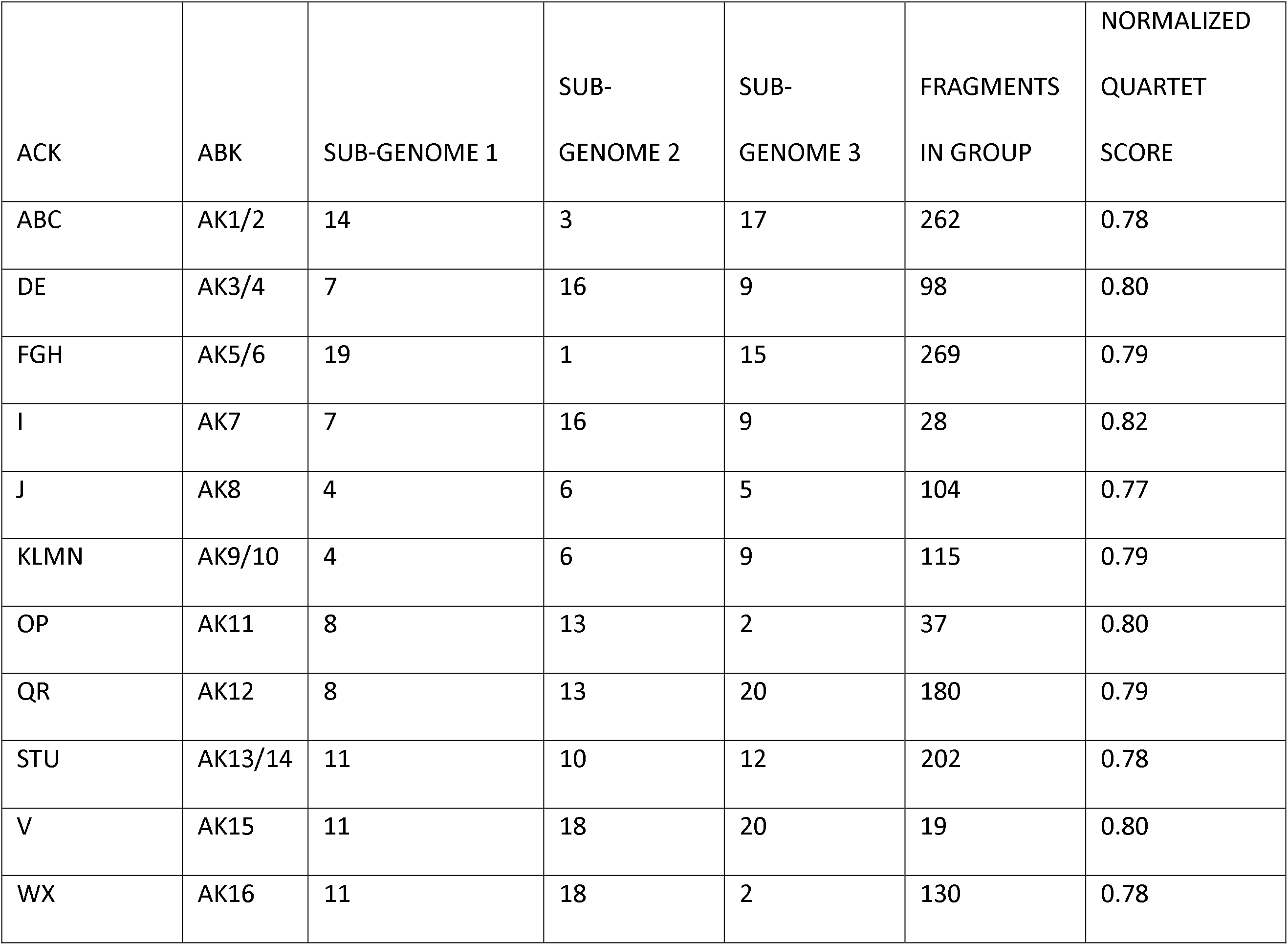
Summary of ACK and ABK groups across *Camelina sativa’s* chromosomes based on bowtie hits showing preservation of ancestral genome structure across ABK regions, but movement of some regions across chromosomes. Chromosomes were assigned to sub-genomes based on summary trees constructed by MrBayes from the ACK fragments grouped by ABK using ASTRAL III. The normalized quartet score, indicating the similarity of trees from individual fragments (out of one), produced by ASTRAL-III is presented. Sub-genome 1 was defined as the genome with closest relationship to *C. neglecta* while sub-genome 3 was defined as that with the closest relationship to the *C. hispida* varieties.

The species trees produced by coalescent modeling by ASTRAL-III from gene trees produced from all fragments within the eleven groups indicated the phylogenetic relationship between *C. sativa*’s chromosomes and the other taxa (Table 4; Table 5; Fig. 1). Trees were largely congruent as indicated by the high normalized quartet scores (Table 4).

**Fig 1.**
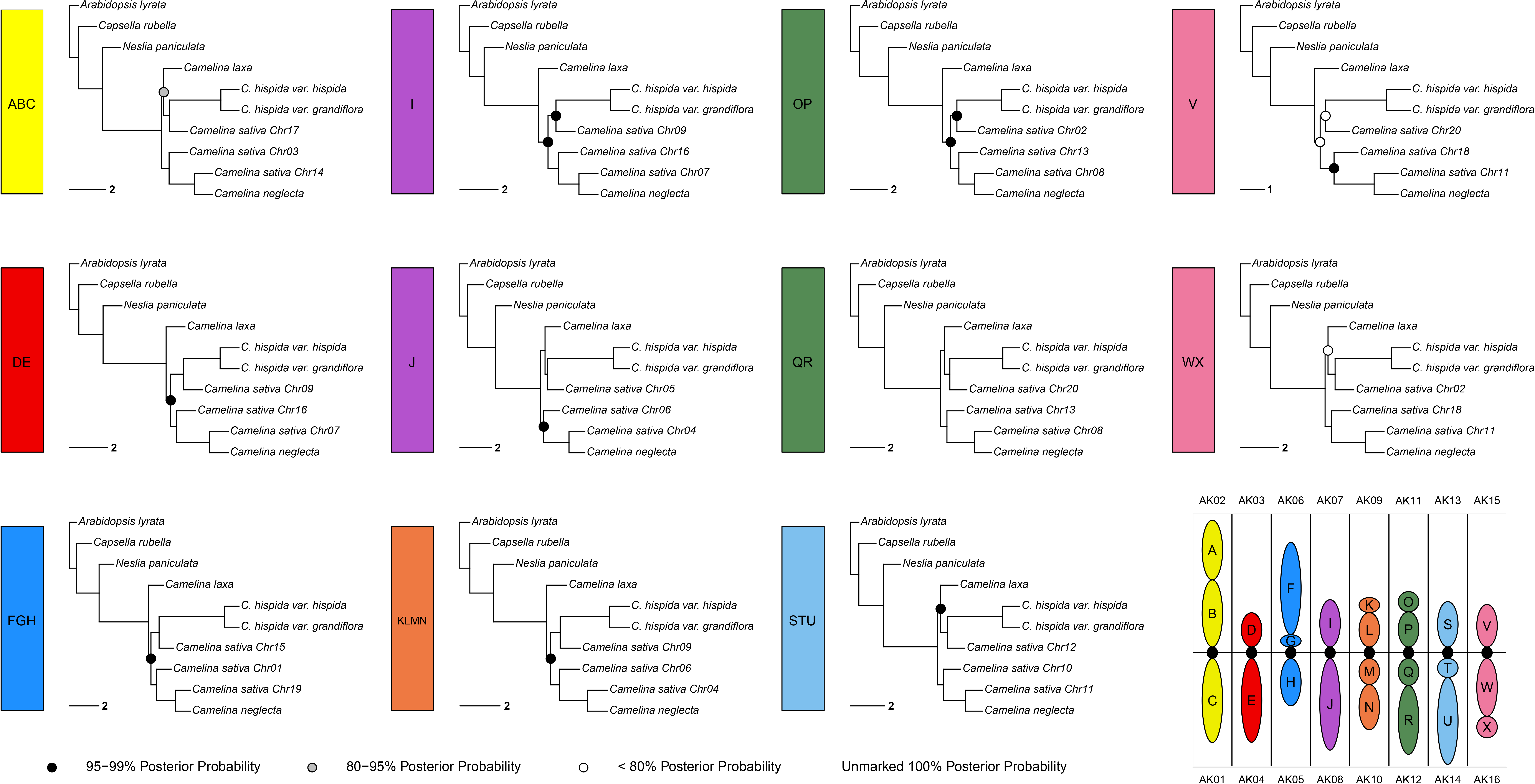
Species Trees. These species trees were produced by ASTRAL-III using a multi-species coalescent model and trees produced by MrBayes for each fragment identified by bowtie 2’s alignment of sequence fragments from *A. lyrata*. These are grouped within eleven ABK groups that correspond to either whole ancestral chromosomes or chromosome arms, that have been largely conserved. The ACK(s) included in each tree are indicated in the box to the left of each tree and the ancestral chromosome structure with division into ACK and ABK is shown in the lower right.

**Table 5.** Sub-genome structure of *C. sativa* based on phylogenetic relationships of AKB regions and diploid genomes determined here in comparison to the original structure proposed by Kagale et al. 2014.

|  | **This Paper** | **Kagale et al 2014.** |
| --- | --- | --- |
| **Sub-genome 1** | Cs 14<br>Cs 07<br>Cs 19<br>Cs 04<br>Cs 08<br>Cs 11 | Cs 17<br>Cs 16<br>Cs 15<br>Cs 04<br>Cs 13<br>Cs 11 |
| **Sub-genome 2** | Cs 03<br>Cs 16<br>Cs 01<br>Cs 06<br>Cs 13<br>Cs 10<br>Cs 18 | Cs 14<br>Cs 07<br>Cs 19<br>Cs 06<br>Cs 08<br>Cs 10<br>Cs 18 |
| **Sub-genome 3** | Cs 17<br>Cs 15<br>Cs 09<br>Cs 05<br>Cs 02<br>Cs 20<br>Cs 12 | Cs 03<br>Cs 05<br>Cs 01<br>Cs 09<br>Cs 20<br>Cs 02<br>Cs 12 |

The consensus species trees estimated by StarBEAST2 using the reciprocal best hit (RBH) sequences also most consistently recovered this topology (Fig. 2 and 3) as did the final species tree predicted by OrthoFinder based 14,513 genes trees (Fig. 4). Further, all methods indicated *Arabidopsis* is the basal genus in this group, while *Neslia* is closest genus to *Camelina* among those studied.

**Fig. 2.**
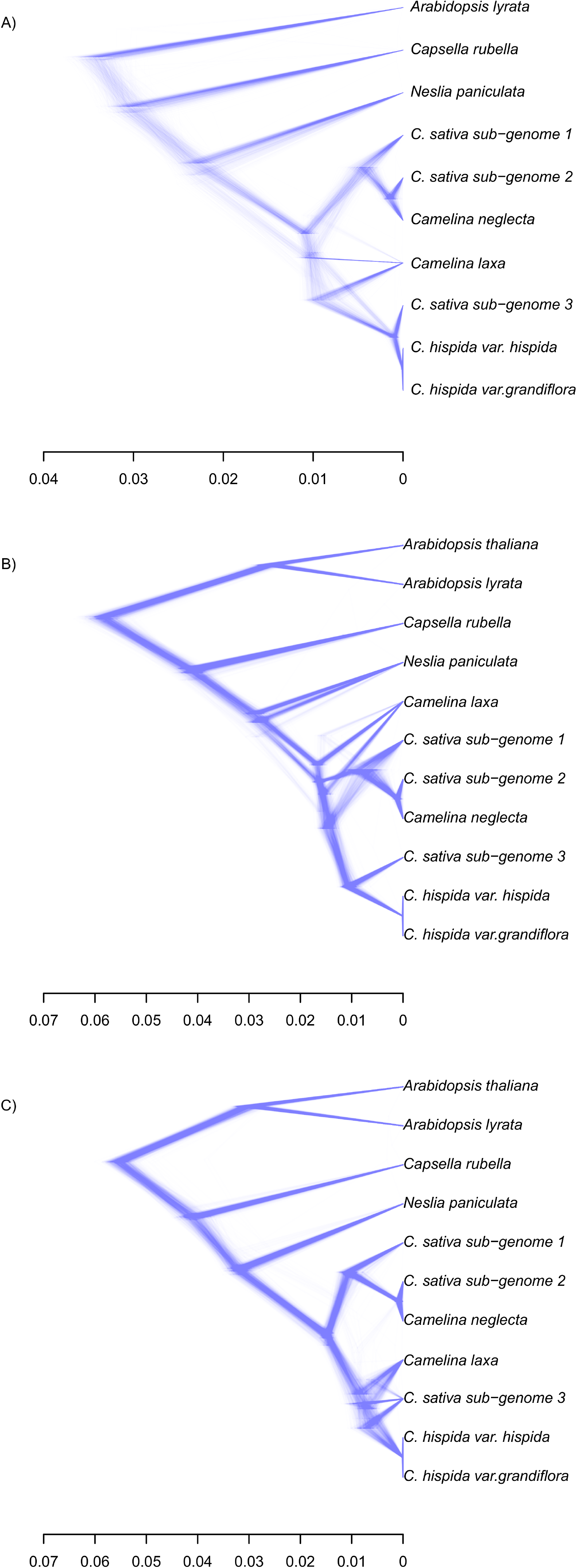
Density tree plots. Density tree plots of 1,000 randomly chosen tree estimated by StarBEAST2 for a set of A) fragment sequences, and B) genes, C) two sets of reciprocal gene sequences.

**Fig. 3.**
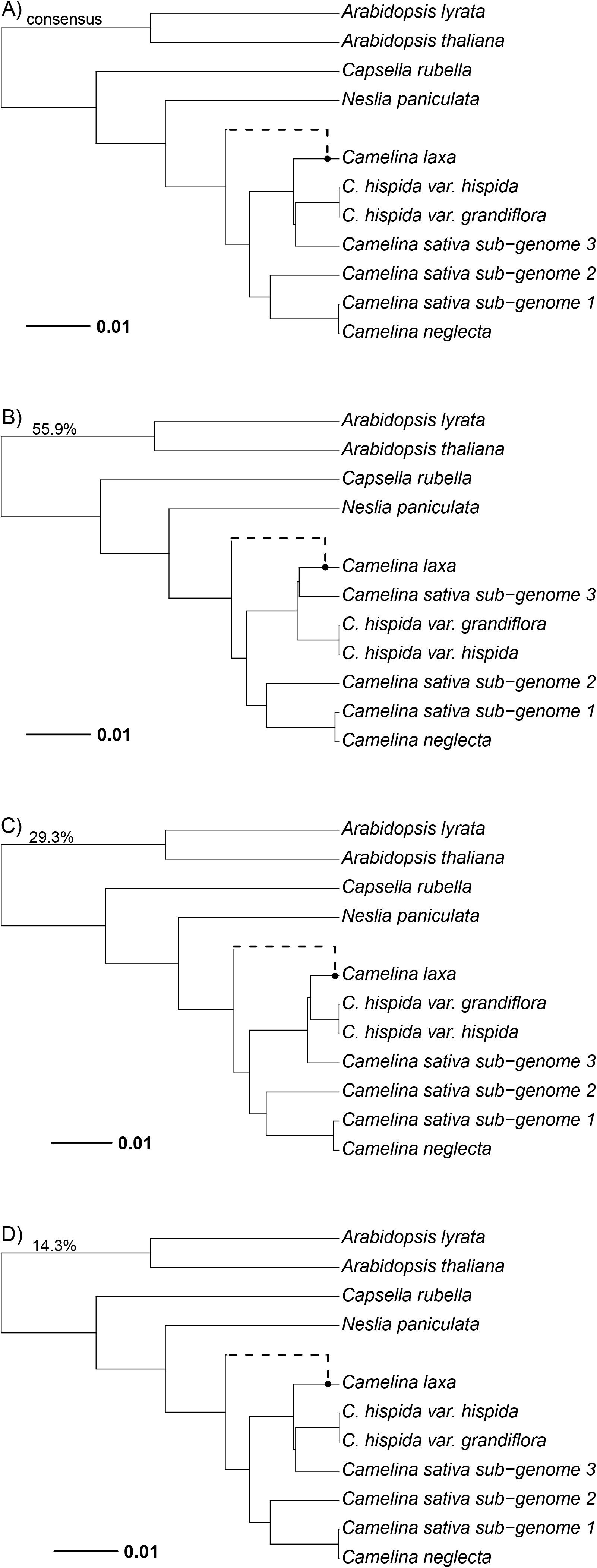
Consensus networks. The consensus networks generated with PhyloNet using one set of reciprocal gene sequences (A) and the three most credible networks contributing to this consensus (B-D) with the percentage of credible networks represented.

**Fig 4.**
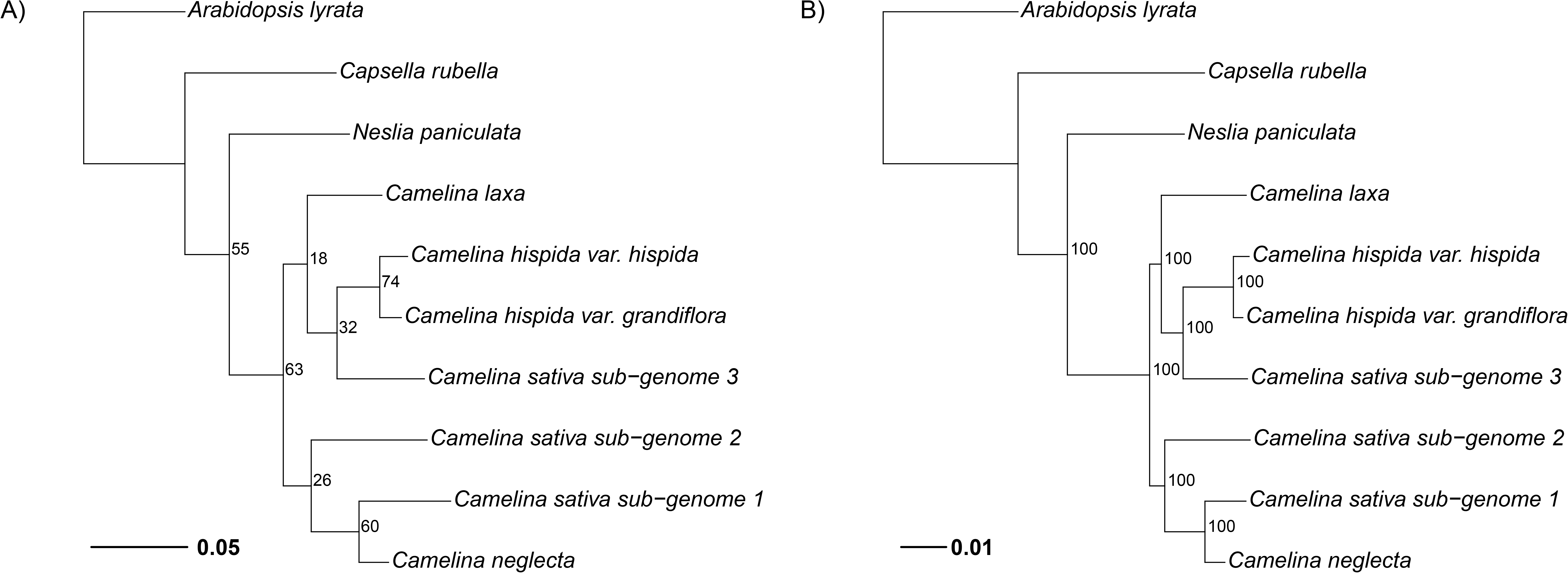
Trees from OrthoFinder. Phylogenetic trees generated by OrthoFinder based on *in silico* predictions of amino acids produced by AUGUSTUS A) a consensus of 14,513 gene trees with an inferred root and B) a concatenated multiple sequence alignment of 7,401 shared, single-copy genes. Node labels indicate support values for each bipartition with the higher levels of support for the concatenated data expected for the type of analysis.

The primary uncertainty in tree topology was the position of *C. laxa* - either basal to all the *Camelina* spp. or in a clade with *C. sativa*’s sub-genome 3 and *C. hispida.* Specifically, of the randomly chosen sets of sequences from across the genome, ASTRAL-III consensus trees produced trees with *C. laxa* basal to the *Camelina* group just over half the time - 14 of 25 for the fragments and 15 of 25 for the reciprocal best hit genes. The consensus trees produced by StarBEAST2 showed greater disparity with 18 of 25 suggesting *C. laxa* is basal for the fragments but only 6 of 25 for the RBH. A thin majority, 14 of 25, of the RBH suggested *C. laxa* in a clade with *C. sativa*’s sub-genome 3. This topology, with *C. laxa* in the sub-genome 3 clade, was also produced by Orthogene (Fig. 4). Density tree plots of trees generated by StarBEAST2 indicated mixed signals for *C. laxa*’s position for sequence fragments (Fig. 2A) and for genes (Fig. 2B). These plots also indicated that some trees differed in the timing of divergence in genes among *C. laxa*, *C. sativa*’s sub-genome 3, and the varieties of *C. hispida* (Fig. 2C). Both ASTRAL-III and StartBeast2 account for incomplete lineage sorting; however, given the propensity of Brassicaceae taxa to hybridize, reticulate evolution between these taxa was investigated with PhyloNet. However, for many of the sets of 24 unlinked fragments or genes, PhyloNet indicated trees without reticulations were the most credible for the fragments (10 of 25) and the reciprocal best hit genes (17 of 25). With other sets one reticulation was included in the credible networks, but their placement was inconsistent (Fig. 3).

### Results for Chloroplasts

The chloroplast (cp) assemblies produced for the four diploid *Camelina* species sequenced here, based on the scaffolding of contigs using *C. sativa*’s cp, resulted in contigs between 152,239 bp (*C. hispida var. grandiflora*) and 153,366 bp (*C. neglecta*) in comparison to the 153,044 bp for *C. sativa’s* cp assembly. Annotation by GeSeq indicated expected features such as the large inverted repeat and that ARAGORN 1.2.38, invoked by GeSeq, found the same 39 genes in all four of the *Camelina*cp’s assembled here and that of *C. sativa.* The cp assemblies varied in the degree of fragmentation of the genes within the assembly with *C. hispida var. grandiflora* showing the greatest number of fragmented features (89) and *C. neglecta* the fewest (20) (Fig. S1). The resulting tree from entire chloroplast sequences of all eleven taxa included in the analysis, showed the chloroplast sequence from *C. sativa* was most closely related to that of *C. neglecta* and that these taxa were sister to a clade containing the varieties of *C. hispida* with *C. laxa* basal. As with the analysis of nuclear genes, the chloroplast indicates *Capsella* as more closely related to *Camelina* than *Arabidopsis* (Fig. 5).

**Fig. 5.**
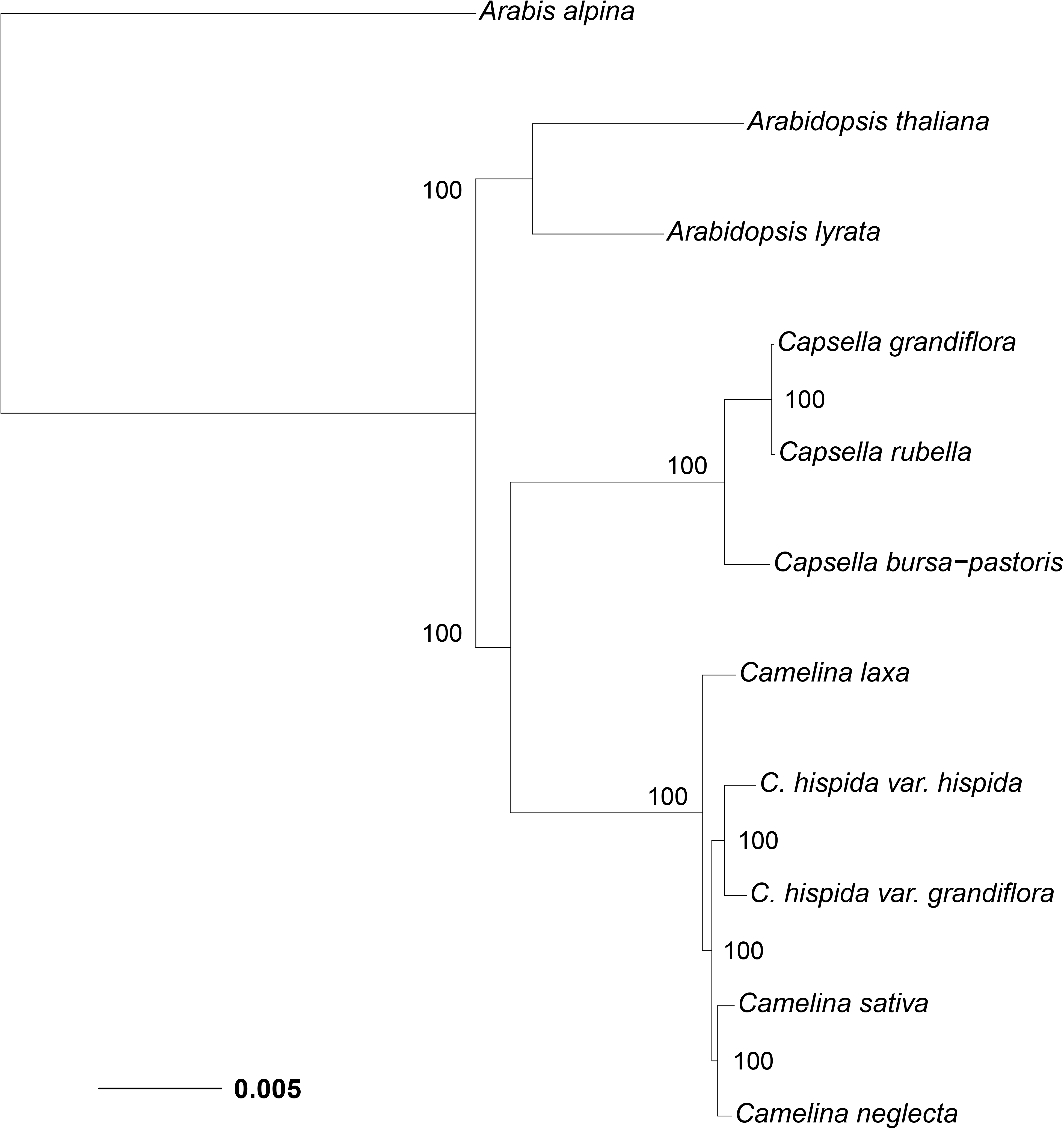
Chloroplast Based Phylogeny. Phylogeny constructed from whole chloroplast alignment using MrBayes. Node labels indicate posterior probabilities.

### Synteny between Camelina diploids and A. lyrata and C. sativa

The four diploid species sequenced here showed extensive synteny with both *A. lyrata* and *C. sativa* genomes with conservation of the ACK and ABK blocks (Fig. 6). The analysis indicates a high level of synteny between *A. lyrata* and *C. neglecta*, and to a greater degree, between *C. neglecta* and *C. sativa*sub-genome 1 (Fig. 6). Similarly, there is strong synteny between *A. lyrata* and *C. hispida* and, to a greater extent, between *C. hispida* and *C. sativa* sub- genome 3 (Fig. 6). In contrast the genome of *C. laxa*has more extensive rearrangements and breaks within ACK and ABK blocks (Fig. 7) including fragmentation of the E block, and separation of parts of the F, J, U, and W blocks on to separate chromosomes. Further, chromosome number reduction in this species involved portions of several of the ancestral chromosomes have being incorporated into several different chromosomes such as the upper arm of chromosome 6 - AK06 (FG) which has been split between chromosomes 3 and 5.

**Fig. 6.**
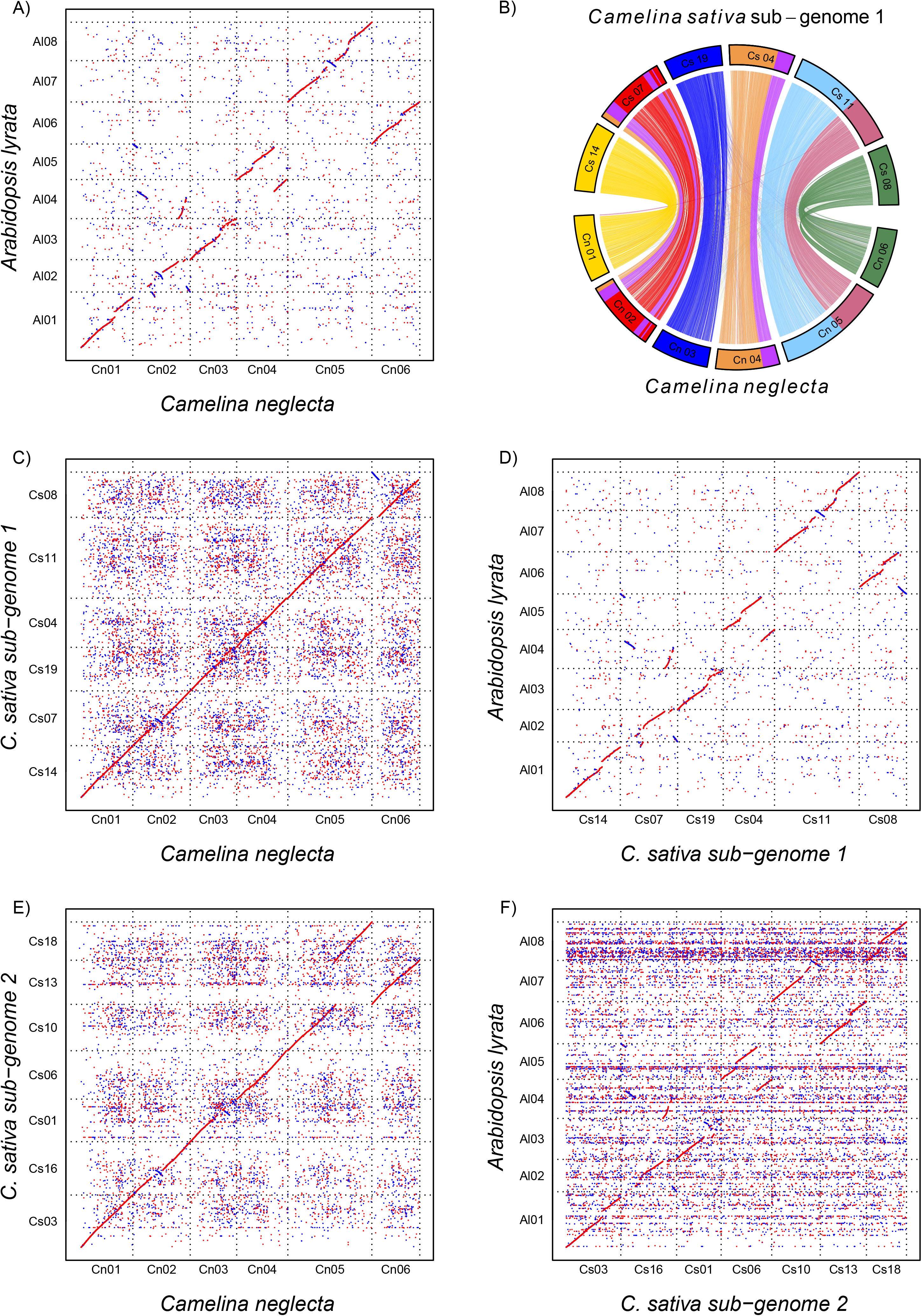
Synteny Plots with *C. neglecta* and *C. sativa’s* sub-genomes 1 and 2. Plots indicating regions of synteny as indicated by nucmer between *C. neglecta*, with *A. lyrata* (A), *C. sativa’s* sub-genomes (B, C, E) and between *A. lyrata* and *C. sativa’s* sub-genomes (D, F). Plot B is coloured by alignment with *A. lyrata’s* chromosomes, analogs for the ancestral chromosomes.

**Fig. 7.**
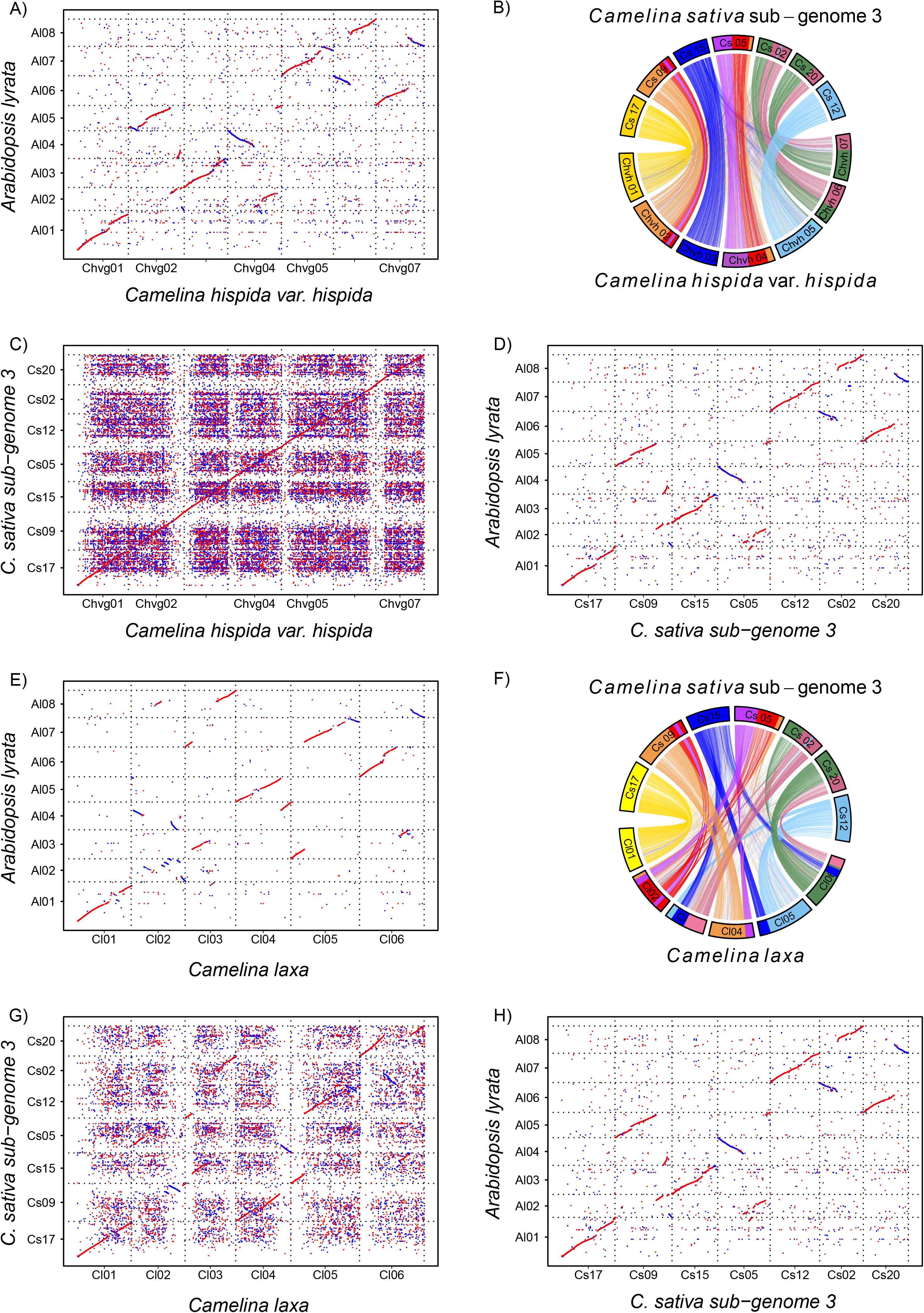
Synteny Plots with *C. hispida* var. *hispida*, *C. laxa* and *C. sativa* sub-genome 3. Plots indicating regions of synteny as indicated by nucmer among the diploid *Camelina* species *C. hispida* var. *hispida* and *C. laxa* with *C. sativa’s* sub-genomes (A, D, G, H) and with A. lyrata (B, E, I). Synteny between *C. sativa* sub-genomes and *A. lyrata* are shown for comparison (C, F). Plot B and F are coloured by alignment with *A. lyrata’s* chromosomes, analogs for the ancestral chromosomes.

### Transposable element annotation and comparison

Annotation of the transposable elements (TEs) by the Extensive *de novo* TE Annotator (EDTA) indicated that TEs made up 34-35% of the diploid *Camelina* genomes, with the exception of *C. hispida* var. *hispida* where TEs accounted for 50% of the genome (Fig. 8, Table S2). The largest groups of TEs identified belonged to the *Helitron* (DHH) and *Gypsy* superfamilies (RLC). EAHelitron identified fewer *Helitrons* than EDTA (Table 6). However, the difference in the number of *Helitrons* in *C. neglecta* and *C. hispida* var. *hispida* was even more divergent with *Helitron* densities of 9.6 and 20.4 per 1,000,000 bp respectively. The percentage of *Camelina hispida* var. *grandiflora*’s genome (34%) and the *Helitron* density (11.8) was much lower than that of *C. hispida* var. *hispida* (Table S2). The three sub-genomes of *C. sativa* also showed differences in the number of TEs and specifically *Helitrons*, with sub-genome 1 and 2 showing similar percentages of TEs at 30.2% and 26.6% and similar *Helitron* densities at 6.1 and 5.8, but sub-genome 3 showing higher values at 40% and 14.7. In syntenic regions, *C.sativa* shared more *Helitrons* with *C. hispida* var. *hispida* (258) than with *C. hispida* var. *grandiflora* (178) (Table 7, Fig 9).

**Fig. 8.**
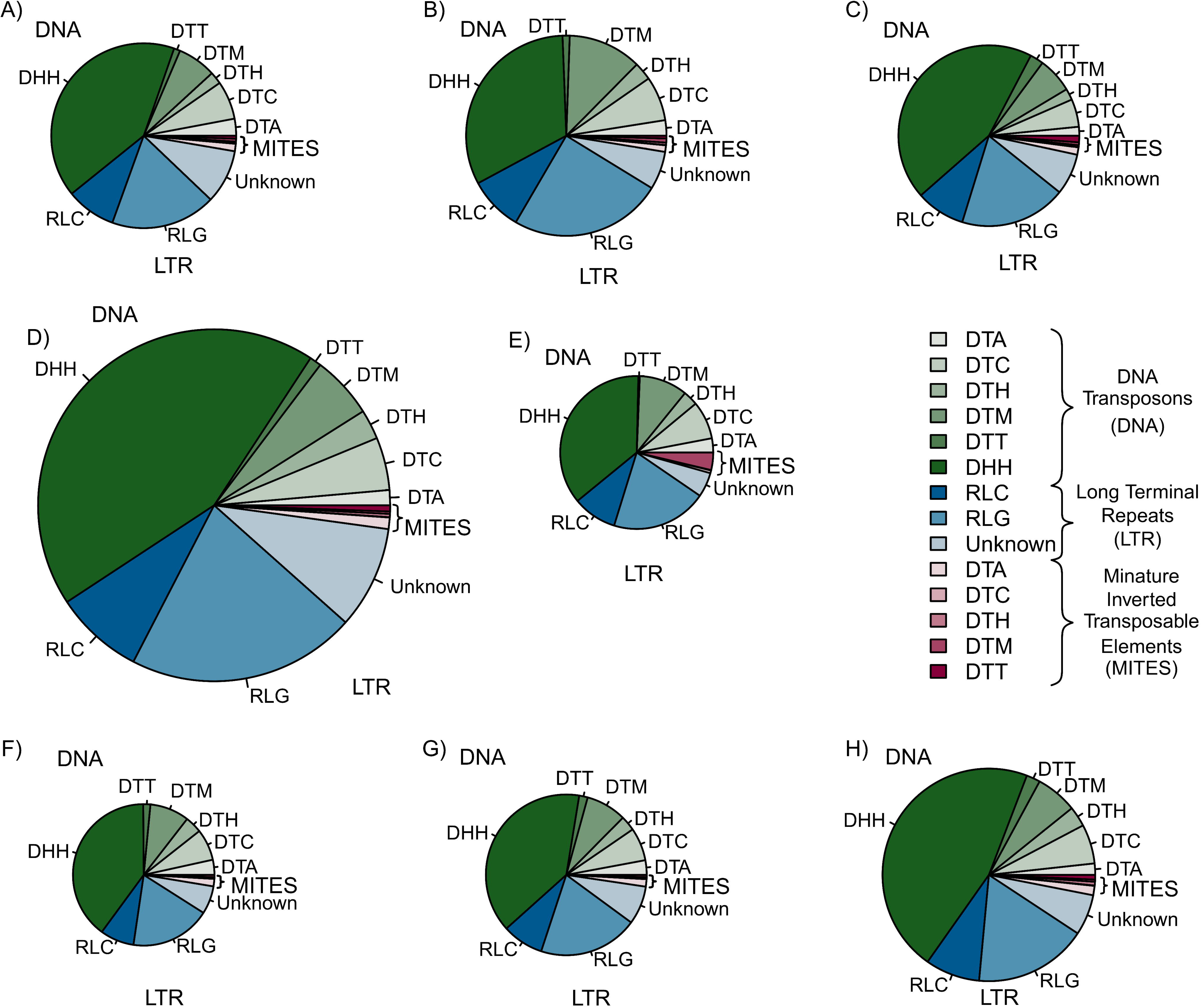
Transposable Element Types and Abundance. Types of transposable elements (TE) in A) *Camelina neglecta*, B) *C. laxa*, C) *C. hispida* var. *grandiflora*, D) *C. hispida* var. *hispida*, E) *Arabidopsis lyrata*, F) *C. sativa* sub-genome 1, G) *C. sativa* sub-genome 2, and H) *C. sativa* sub-genome 3, as annotated by Extensive De novo Transposable Element (EDTA). Pie graphs are scaled by the percentage of the genome attributed to TEs which at highest is 49.6% for *C. hispida* var. *hispida* (D). Classification of transposable element type follows the unified classification system for eukaryotic transposable elements that uses a three letter code indicating class, order and superfamily (34). Here these are divided into three groups: DNA transposons (DNA), which includes *Helitrons* (DHH); long terminal repeats (LTR), which includes retrotransposons RLC (*Copia*) and RLG (*Gypsy*); and miniature inverted transposable elements (MITES).

**Fig. 9.**
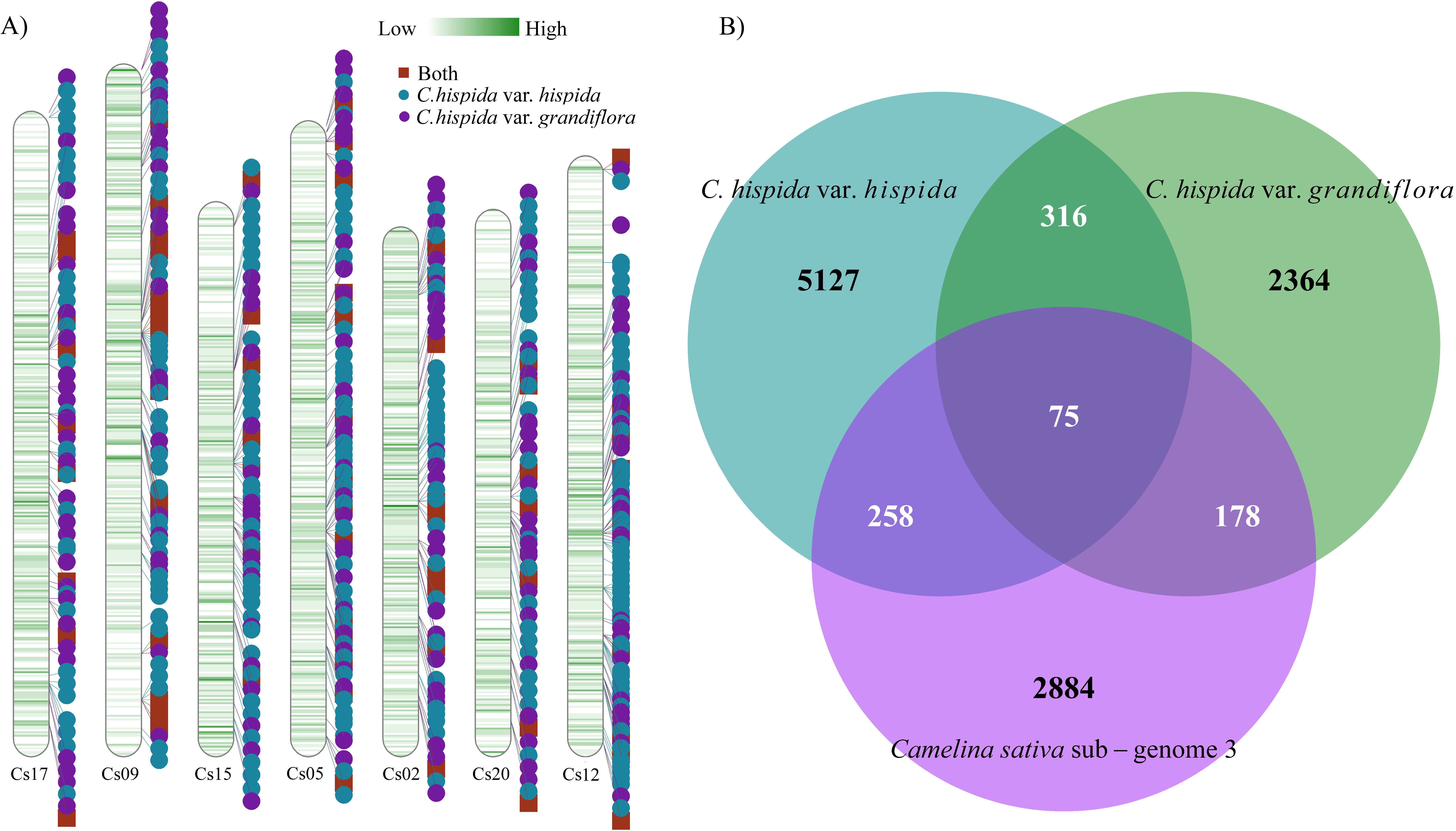
Helitron Ideogram and Abundance. A) Density of *Helitrons* identified by EAHelitron across the sub-genome 3 of *C. sativa* with the locations of *Helitrons* shared by *C. hispida* var. *hispida*, *C. hispida* var. *grandiflora* or both indicated and B) Venn diagram of the number of Helitrons shared among *C. hispida* var. hispida, *C. hispida* var. *grandiflora*, and *C. sativa’s* sub-genome 3.

**Table 6.**
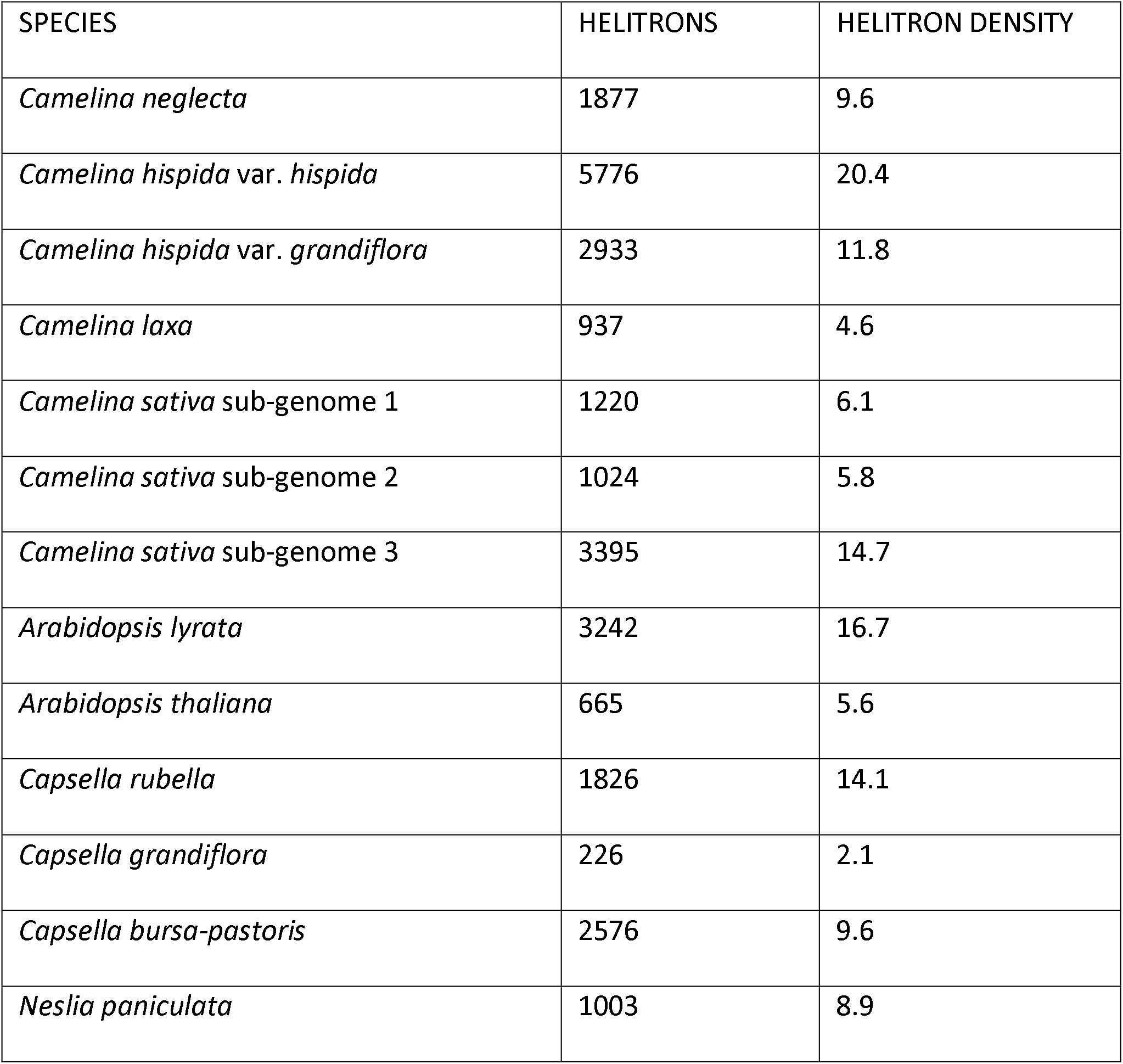
Results of *Helitron* (DHH) annotation by EAHelitron. *Helitron* density is calculated as the number of *Helitrons* detected in the genome for each 1,000,000 bp.

| SPECIES | HELITRONS | HELITRON DENSITY |
| --- | --- | --- |
| *Camelina neglecta* | 1877 | 9.6 |
| *Camelina hispida* var. *hispida* | 5776 | 20.4 |
| *Camelina hispida* var. *grandiflora* | 2933 | 11.8 |
| *Camelina laxa* | 937 | 4.6 |
| *Camelina sativa* sub-genome 1 | 1220 | 6.1 |
| *Camelina sativa* sub-genome 2 | 1024 | 5.8 |
| *Camelina sativa* sub-genome 3 | 3395 | 14.7 |
| *Arabidopsis lyrata* | 3242 | 16.7 |
| *Arabidopsis thaliana* | 665 | 5.6 |
| *Capsella rubella* | 1826 | 14.1 |
| *Capsella grandiflora* | 226 | 2.1 |
| *Capsella bursa-pastoris* | 2576 | 9.6 |
| *Neslia paniculata* | 1003 | 8.9 |

**Table 7.** The number of shared Helitrons detected by EAHelitron in *C. sativa’s* sub-genome 3 within regions of synteny among *C. sativa’s* sub-genome 3, *C. hispida* var. *grandiflora*, and *C. hispida* var. *hispida* chromosome.

| *C. sativa*<br>CHROMOSOME | *C. sativa*<br>HELITRON<br>COUNT | HELITRONS SHARED WITH |  | COMMON<br>HELITRONS |
| --- | --- | --- | --- | --- |
|  |  | *C. hispida* var. *grandiflora* | *C. hispida* var. *hispida* |  |
| Cs 17 | 516 | 24 | 31 | 9 |
| Cs 09 | 518 | 29 | 41 | 17 |
| Cs 15 | 456 | 27 | 38 | 7 |
| Cs 05 | 513 | 27 | 59 | 14 |
| Cs 12 | 505 | 21 | 35 | 12 |
| Cs 02 | 496 | 27 | 29 | 9 |
| Cs 20 | 391 | 23 | 25 | 7 |
| Total | 3395 | 178 | 258 | 75 |

## Discussion

Whole genome sequencing is allowing a deeper understanding of the processes that have shaped plant evolution including polyploidization, hybridization and chromosomal rearrangements. Allopolyploid crops have become excellent systems for understanding these genomic changes because they have often received substantial sequencing effort (e.g. *Triticum aestivum* L. (Appels et al. 2018), *Solanum lycopersicum* L. (Sato et al. 2012), *Zea mays* ssp. *mays* L. (Schnable et al. 2009; Jiao et al. 2017), and *Arachis hypogaea* L. (Bertioli et al. 2019)). Moreover, as work on these species has shown, a greater understanding of the genomic changes associated with allopolyploidization or autopolyploidization, the genomic consequences of domestication, and the potential breeding resources can be achieved when related, extant diploid species are also sequenced (e.g. Sato et al. 2012; Marcussen et al. 2014; Bertioli et al. 2016; Ramos-Madrigal et al. 2016).

The genome of the allohexaploid *Camelina sativa* (L.) Crantz (camelina; 2n = 40) has been sequenced (Kagale et al. 2014), but extant relatives of the three parental genomes have not. Four diploid taxa are known within the genus: *Camelina neglecta*, *Camelina laxa*, *Camelina hispida* var. *hispida* and *Camelina hispida* var*. grandiflora*. Here we sequenced the genomes of these four diploids, assembled high quality chromosome level drafts, and examined their phylogenetic relationship with *C. sativa’s* sub-genomes. Each genome showed synteny with *A. lyrata* with conservation of ACK and ABK blocks, with each region mapping once, confirming a diploid structure (Fig. 6, 7). This conservation of ACK and ABK blocks allowed for further dissection of the genomes and phylogenetic analysis.

### Camelia sativa’s Sub-genomes

Species trees produced by ASTRAL-III’s multi-species coalescent model using trees produced from sequence data from each ABK or ACK indicated the phylogenetic relationships between these regions from *C. sativa*’s chromosomes and the other taxa included (Fig. 1). These phylogenies indicate that the phasing of *C. sativa*’s chromosomes to sub-genomes required reassessment compared to that originally suggested by Kagale et. al ( 2014). Given that rearrangement and fractionation have not been predominant mechanisms of genome change in *C. sativa* (Lysak et al. 2016), limited visual information is available for this division. Phasing polyploid genomes into their sub-genomes remains a challenging issue (Rothfels 2021) and the majority of tools to aid in this are currently only able to handle tetraploids (e.g. AlloppNET) (Jones 2013). Here, phasing was accomplished by using the distribution of ancestral syntenic blocks and placing them in phylogenetic context with related diploids. This allowed us to define sub-genome 1 as composed of the chromosomes with the closest phylogenetic relationship with *C. neglecta*, sub-genome 2 as those chromosomes sister to the clade with *C. neglecta* and sub- genome 1, and sub-genome 3 as chromosomes with the closest phylogenetic relationship to the varieties of *C. hispida* (Table 4; Table 5; Fig. 1). This is a different nomenclature than our previous work which defined sub-genome 2 as most closely related to *C. neglecta* (Lujan Toro 2017), but it is concordant with the revised sub-genome definition recently published for *C. sativa* (Chaudhary et al. 2020).

In Chaudhary et al.’s analysis (2020), the authors characterized 193 accessions of *Camelina,* including *C. neglecta*, *C. laxa*, *C. hispida*, *C. rumelica* Velen., tetraploid and hexaploid *C. microcarpa* Andrz. ex DC., and *C. sativa*, using a genotyping by sequencing (GBS) approach that mapped SNPs in sequences for these accessions to *C. sativa*’s genome. They determined that sequences from *C. neglecta* aligned to six of *C. sativa*’s chromosomes and tetraploid *C. microcarpa*aligned with 13 of the chromosomes. However, they determined that alignment reads did not correspond to the sub-genome structure published in Kagale et. al (2014) and refined the composition of the sub-genomes accordingly with *C. neglecta* sequences aligning to sub-genome 1, the tetraploid *C. microcarpa* sequences aligning to sub-genomes 1 and 2, and *C. hispida* partially aligning to sub-genome 3 (Chaudhary et al. 2020). As a result very different approaches, an analysis based on SNPs and a phylogenetic analysis of whole genome sequences, has resulted in the same conclusions about refinements to the sub-genome structure of *C. sativa*.

In all three sub-genomes, the majority of the changes in the genome structure in comparison to *A. lyrata* are shared between the diploid genomes and the sub-genomes of *C. sativa* (see below, Fig. 6, 7). This conservation of structure in these closely related genomes is in line with Lysak et al.’s expectations that the majority of the changes in chromosome structure seen in *C. sativa* compared to the ancestral chromosomes were likely present in the ancestral diploid genomes (Lysak et al. 2016). While the formation of *C. sativa* was relatively recent (Kagale et al. 2014), it is clear that allopolyploidization did not result in extensive structural reorganization of the genome, but rather can be characterized by conservation of ancestral chromosome structure and gene order.

Inter-genome recombination has been observed in some allopolyploids, including resynthesized *Brassica napus* (Pires et al. 2004), and if this has occurred in *C. sativa*’s genome it could result in conflicting signals in some loci, as could historical gene flow and introgression. Some genes appear to show a signal of hybridizations among *C. laxa*, *C. sativa*’s sub-genome 3 and the varieties of *C. hispida* (Fig 2., Fig 3.), and further evaluations of this possibility could be completed with broader, population level sampling of the taxa and evaluation of the evidence of introgression (Martin et al. 2015; Martin and Jiggins 2017; Crowl et al. 2020).

### *Camelina neglecta* - an extant, maternal relative

With the refinements to the phasing of the sub-genome structure of *C. sativa*, in addition to the close phylogenetic relationship, there is strong synteny between the draft *C. neglecta* and *C. sativa*’s sub-genome 1 (Fig. 6) with similar conservation of the ancestral chromosome structure. Specifically, only two large inversions are apparent when *C. neglecta*’s genome and *C. sativa*’s sub-genome 1 are compared: one at the start of *C. neglecta’*s chromosome 6 and end of *C. sativa’s* chromosome 8 and one within *C. neglecta’*s chromosome 2 and *C. sativa’s* chromosome 7, which is echoed in the homologous chromosome of *C. sativa*’s sub-genome 2. Interestingly, the inversion in *C. neglecta’*s chromosome 6 is more similar *A. lyrata*’s chromosome 6 suggesting this change may have occurred in *C. sativa*’s genome 1 following allopolyploidization. This similarity and the chloroplast data, which suggests the maternal lineage of *C. sativa* is most closely related to *C. neglecta,* indicates that *C. neglecta* is a close extant, maternal relative of *C. sativa*’s sub-genome 1.

*C. neglecta* is also the closest relative of sub-genome 2 (Fig. 6) with the largest difference in structure the lack of fusion in *C. sativa*’s chromosomes 10 and 18 compared to *C. neglecta’*s chromosome 5 and several additional areas of inversions within *C. neglecta’*s chromosome 3. This suggests that the taxon that contributed *C. sativa*’s sub-genome 2 is or was a close relative of *C. neglecta* that retained the ancestral chromosome structure of n = 7 reconstructed for the genus by Mandáková et al. (2019). This taxon is currently unknown, but given that *C. neglecta* was only recently described as a distinct species because of its morphological similarity to *C. microcarpa*, it is possible that the taxon has been collected, but is similarly cryptic and not yet identified as distinct from accessions of *C. microcarpa*.

### *Camelina hispida* - an extant, paternal relative

The varieties of *C. hispida* show a close phylogenetic relationship with *C. sativa’s* sub- genome 3 (Fig 7). There is extensive synteny among these genomes indicating strong preservation of chromosome structure and gene order (e.g. Fig. 7). As in the comparisons between *C. neglecta* and *C. sativa*’s genomes 1 and 2, three chromosomes appear to have inversions, one of which is shared in comparisons with *A. lyrata* and potentially represents a change since *C. sativa’s* formation. While the phylogenies indicate that both varieties of *C. hispida* share a common ancestor with *C. sativa*’s sub-genome 3, the frequency of TE elements differs strongly between the two varieties and is responsible for the 12% larger 2C DNA content of in *C. hispida* var. *hispida* (Martin et al. 2017).

The frequency of TEs and, in particular, shared *Helitrons* suggest *C. hispida* var. *hispida*’s genome may more closely resembles the genome that contributed *C. sativa’s* sub- genome 3. The frequency of TEs varies across *C. sativa*’s sub-genome’s with sub-genome 3 having the highest number of these elements, at 40% of the sub-genome, compared to 30% in the other two (Fig. 8., Table 6, SI Table 2). Among the diploids sequenced here this percentage is closest to the percentage observed for *C. hispida* var. *hispida* (50%). The largest portion of these TEs by count, *Helitrons*, make up 15% of *C. sativa*’s sub-genome 3 and 16% of *C. hispida* var. *hispida’s* genome. *Helitrons* were first discovered in the genomes of *Arabidopsis thaliana*, *Oryza sativa* L. and *Caenorhabditis elegans* Maupas (Kapitonov and Jurka 2001) and are the most abundant TEs in *A. thaliana*’s genome. They make up the second largest TE portion of the genome after the less numerous, but longer retro-elements such as *Copia* and *Gypsy* (Quesneville 2020). Hu et al. (2019) have suggested that, because *Helitron* density is highly evolutionarily labile they could be used for species identification. In this case, we investigated whether they may provide clues to which genome was more similar to that which contributed to the formation of *C. sativa*. We used the software developed by Hu et al. (2019), EAHelitron, to determine whether *Helitrons* detected in *C. sativa*’s sub-genome 3 were more often shared with *C. hispida* var. *hispida* or *C. hispida* var. *grandiflora*. This indicated that 7.5% of the *Helitrons* in *C. sativa*’s sub-genome 3 are shared with *C. hispida* var. *hispida,* while 5.2% are shared with *C. hispida* var. *grandiflora’s* (Table 7, Fig. 9). This suggests that the genome that contributed *C. sativa*’s sub-genome 3 may have been was more similar to the genome of *C. hispida* var. *hispida* with its higher TE content. However, it is also possible that with the large expansion of TEs in *C. hispida* var. *hispida’s* genome, shared *Helitrons* were more likely to be retained.

Here we see reduced TEs abundance and *Helitron* density in sub-genome 3 from levels observed in *C. hispida* var. *hispida* and in sub-genome 1 compared to *C. neglecta*. However, one expectation is that genomic shock and relaxed selection pressure following allopolyploidization allows an increase in TE abundance (Parisod and Senerchia 2012; Vicient and Casacuberta 2017). For example, proliferation of TEs has been observed in allotetraploid *Coffea arabica* L.. Specifically, *copia* elements have increased the size of sub-genome C compared to that observed in the diploid representative, *C. canephora* L., a representative of the progenitor genome (Yu et al. 2011). Similarly, the transposon *Sunfish*, was observed to be released from repression in the early generations of synthetic autopolyploids of *Arabidopsis thaliana* and *A. arenosa* (L.) Lawalrée as well as their allopolyploid derivative *A. suecica* (Fries.) Norrlin. However, the same was not true for other TE families in *A. suecica* (Madlung et al. 2005) and more recent work by Burns et al. (2021) concluded there was no upregulation in transposon activity in *A. suecica* compared to its ancestral species. Ågren et al. (2016) also determined that the allotetraploid *Capsella bursa-pastoris* did not show genome wide TE proliferation compared to its progenitors, though did note a higher abundance in retrotransposons in gene rich regions, and Sarilar et al. (2013) determined that early generations of synthetic allotetraploid *Brassica napus,* did not show increased activity in three groups of TEs. Interestingly, similar to our observation, Hu et al. (2019) observed that *Helitron* density was also lower in the sub-genomes of the allotetraploid *Brassica napus* L. compared to the extant representatives of the parental genomes *B. rapa* L. and *B. oleraceae* L.. Here, the repeatome of *C. sativa* is less complete than that of the diploid genomes assembled with long reads and this may mean that the *Helitrons* in *C. sativa*’s genome are underrepresented. Alternatively, the diploid lineages may have seen proliferation of TEs since *C. sativa’s* formation. As in the systems explored above, resynthesis of *C. sativa,* could determine whether the apparent changes are repeatable. Further, the variation in TE abundance between *C. hispida* varieties and the presence of extant G1G3, *C. rumelica,* and G1G2, “*C. microcarpa* 4x”, tetraploids (Mandáková et al. 2019; Chaudhary et al. 2020) in addition to the G1G2G3 hexaploid *C. sativa* provides an intriguing opportunity to use resynthesis to study TE dynamics after allopolyploidization and investigate how if TE abundance contributes to sub- genome dominance (Bird et al. 2018; Alger and Edger 2020).

The results presented by Žerdoner Čalasan et al. (Žerdoner Čalasan et al. 2019) suggest that *C. anomala*, a taxon with unknown chromosome number, is sister to *C. hispida* raising the possibility that this species could also be a close relative of *C. sativa*’s sub-genome 3. However, the species has not been collected since the 1800s and remains largely a mystery.

### Camelina laxa

*Camelina laxa* is morphologically distinct from the other taxa studied here with its flexuous or zig-zagging stems and also has the most diverged genome from the ancestral karyotype. It is clear from the phylogenetic evidence here and cytological evidence presented by Mandáková et al. (2019) that *C. laxa* and *C. neglecta* represent separate transitions from n = 7 to n = 6. In *C. laxa,* extensive chromosomal rearrangements occurred, resulting from chromosome shattering, while in *C. neglecta*chromosome reduction resulted from the fusion of two chromosomes (Mandáková et al. 2019). However, the position of *C. laxa* in the phylogenetic tree was the least stable element across all analyses with two main alternatives, either as basal to all other *Camelina* species sampled or as basal to the clade containing the varieties of *C. hispida* and *C. sativa*’s sub-genome 3. While Mandáková et al.’s (2019) analysis of 48 single copy genes using ASTRAL placed *C. laxa* in a clade with *C. hispida*, the consensus from previous work is that *C. laxa* should be considered basal to the other species of *Camelina*. For example, Brock et. al. (2018)’s maximum likelihood consensus phylogeny constructed from ddRADseq for 48 specimens from gene bank material and field collections of *C. sativa*, *C. microcarpa*hexaploids, *C. rumelica*, *C. laxa* and *C. hispida* placed *C. laxa* was basal to the other species. Similarly, a neighbor joining tree produced using GBS data indicated *C. laxa* was basal except the G1G3 tetraploid *C. rumelica*(Chaudhary et al. 2020). Finally, in a more comprehensive analysis of the Camelineae, which included *C. alyssum* (Mill.) Thell. and *C. anomala* Boiss. & Hausskn. ex Boiss. as well as representatives from many other genera included in the tribe *Nelisa, Capsella, Arabidopsis, Catolobus, Pseudoarabidopsis,* and *Chrysochamela*, Žerdoner Čalasan et al. (2019) also placed *C. laxa* as basal to *Camelina*. As a result, the preponderance of evidence currently suggests that this, more diverged lineage, is basal to the *Camelina*.

## Conclusion

As domestication has decreased genetic variability in *C. sativa*accessions (Manca et al. 2013) other members of the genus might be of value for the improvement of *C. sativa*. Here our results indicate that *C. neglecta* and *C. hispida* var. *hispida* could be useful species to investigate variation in traits such as seed size, disease resistance and oil profiles. However, the collections available for these species are limited and international efforts to collect and preserve a broader selection of germplasm for the genus should be considered as we look to these species as sources of desirable traits for the improvement of the crop species (Ford-Lloyd et al. 2011).

## Data Availability

All assembled genomes are available from the National Center for Biotechnology Information as part of Bioproject PRJNA750147.

## Acknowledgements

We would like to thank the ORDC growth facility team for support rearing the plant material, the ORDC Molecular Technology Laboratory for support with sequencing, and the U.S. National Plant Germplasm System and GRIN-Global for conserving and providing the germplasm needed for this research. We thank Dr. S. Wright for sharing the draft genome sequence of *Neslia paniculata* and Julia Mata for help processing convergence data. Funding was provided by Agriculture and Agri-Food Canada as part of the project “Gene flow, diversity and relationships within the Brassicaceae: Focus on *Camelina*” (J-001029).

