## Supplementary Figure Text for "Insights from the genomes of four diploid *Camelina* spp."

**Fig.** **S1** Chloroplasts for the four diploid *Camelina* species annotated by GeSeq and drawn by OrganellarGenomeDraw available through CHLOROBOX (see Materials and Methods). Genes noted on the inside of the chloroplast’s ring are transcribed clockwise while those on the outside are transcribed counter-clockwise. Genes with introns are marked with an asterisk and the inner grey ring indicates GC content with a line indicating 50%. The abbreviation LSC indicates the large single copy region and the short single copy region is labelled SSC.
