## Supplementary figures and images for "Insights from the genomes of four diploid *Camelina* spp."

### Supplementary Fig 1

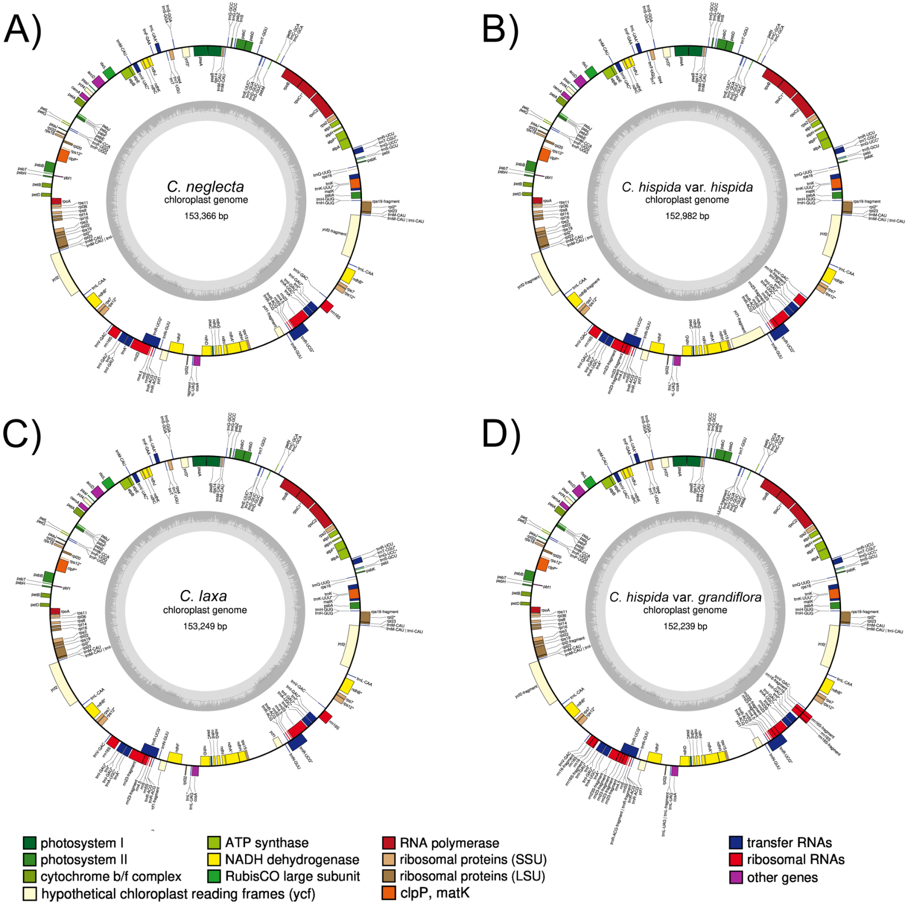
